# Focal adhesion prevention on nanoparticle substrates upregulates stemness gene expression in primary melanocytes

**DOI:** 10.64898/2025.12.28.693089

**Authors:** Benjamin C. U. Tai, Siti Thaharah Mohamed, Jia Kai Lim, Nina K. L. Ma, Andrew C. A. Wan

## Abstract

**Hypothesis:** Cell focal adhesions are initiated by interactions of cell surface integrins with extracellular matrix (ECM) proteins, and cells which lack focal adhesions have been associated with an intermediate adhesive state, which is more labile towards cell reprogramming. We hypothesized that cells cultured under conditions where they exhibit reduced focal adhesions will assume an intermediate adhesive state with a more stem cell-like transcriptional profile.

**Experiments:** To recreate culture conditions for cells to exhibit reduced focal adhesions, we synthesized PMMA nanoparticles by evaporation-induced self-assembly (EISA), employing varying conditions of surfactant concentrations, and allowed them to self-assemble into nanoparticle substrates. Following guidelines from previous work, we selected nanoparticle substrates of dimensions ∼70nm, suitable for the prevention of focal adhesion formation. Primary human melanocytes were cultured on these substrates to verify the reduction of their focal adhesions; following that the cultures were characterized in terms of their expression of distinct sets of genes related to stemness and differentiation.

**Findings:** The melanocytes showed reduced focal adhesions on both non-coated and fibronectin-coated nanoparticle substrates, when compared to melanocytes cultured on TCPs. Accompanying the reduction in focal adhesions, the cells exhibited an increase in the expression of general as well as melanocyte-specific stemness related genes such as OCT4, LIN28, NANOG and PAX6 and a concurrent downregulation of genes expressed by differentiated melanocytes, such as TYR, TRP1 and MITF. These observations support our hypothesis.

## Introduction

Focal adhesions are large, multi-protein structures through which the cellular cytoskeleton connects to the extracellular matrix (ECM)[1]. In response to cell adhesion to the ECM, mechanical forces and biochemical signals are transmitted through the focal adhesions to affect the gene expression, cell differentiation and proliferation, and ultimately the survival of the cell. *In vivo*, a class of proteins called the matricellular proteins, including SPARC, tenascin C and thrombospondin, are able to induce the focal adhesion disassembly, leading to an intermediate state of cell adhesion. Such an adhesive state has been proposed to be involved in tissue processes such as remodelling and wound healing, and involve a differential gene expression profile[2, 3].

From previous work, we have found that ECM signalling is dependent on the dimensional context of the cell[4]. Glioma cells that were exposed to specific laminin isoforms in a 3D electrospun scaffold exhibited an enhanced stemness phenotype compared to cells that were grown on 2D tissue culture plates. Following that, we hypothesized that cells that lack focal adhesions would respond to ECM signalling in a different manner from cells that are strongly adhered, with respect to their expression of genes involved in stemness and differentiation.

Instead of treating the cells with matricellular proteins (which interact with cells and ECM in other ways and thus present confounding effects), we have employed an alternative, physical approach to disrupt focal adhesion assembly in the present work. Culture of cells on substrates of various nanotopographies have shown that nanostructural features of specific dimensions prevent cells from forming focal adhesions[5]. To create such nanostructural features, we synthesized submicron sized PMMA particles and allowed them to self-assemble into planar nanoparticle substrates. We selected the substrate type that met the dimensional criteria for focal adhesion prevention, for subsequent cell culture experiments. Primary human melanocytes were cultured on these substrates to verify the reduction of their focal adhesions; following that the cultures were characterized in terms of their expression of distinct sets of genes related to stemness and differentiation.

## Materials and Methods

### Fabrication of PMMA nanoparticle substrates

#### Preparation of polymethyl methacrylate (PMMA) polymeric nanoparticle solution

PMMA nanoparticles were synthesized via a differential microemulsion polymerization technique[6, 7]. This reaction was carried out in the presence of a surfactant sodium dodecyl sulfate (SDS, Promega) and catalyzed by an initiator, ammonium persulfate (APS, Promega). Briefly, 8.3 ml of methyl methacrylate (MMA, Sigma Aldrich) monomer was added drop wise using a syringe pump at a rate of 0.15 ml/min, to a solution containing 47.6 mg of APS and 5 mg of SDS. This gave a final surfactant concentration of 0.60 mg/mL. The reaction was mixed homogenously using a stirrer bar and maintained at 75° C. The broth was left to stir for an additional 1 hour for the solution to age. In a separate synthesis, 52.9 mg of SDS was added, corresponding to a higher final surfactant concentration of 6.4 mg/mL.

#### Preparation of substrates

2 mL of the nanoparticle suspension was placed on ice, to which 500 uL of 1.5% P-123 and 1 mL of cold ethanol was added, followed by vortexing. The nanoparticle suspensions were poured into on a non-adhesive aluminium mould of diameter 2.3 cm and allowed to stand in the oven at 30 ºC. The moulds were left for evaporative induced self-assembly (EISA) to occur for 72 hours, resulting in closely packed arrays of nanoparticles. The compacted substrates were then sintered in a pre-heated furnace maintained at 120 ºC for about an hour.

#### Primary melanocyte cell culture

Normal human primary epidermal melanocytes (PEM) from neonatal foreskin was obtained from American Tissue Culture Collection (ATCC, Manassas, VA, USA) and maintained in dermal cell basal medium (ATCC, USA) supplemented with melanocyte growth kit (ATCC, USA) and penicillin-streptomycin-amphotericin B (Gibco, USA). The cells were incubated at 37 °C with 5% CO_2_. In the present work, we used melanocytes which had been passaged less than 10 times and on average, 8 times. Prior to cell seeding on the PMMA nanoparticle substrates, the substrates were sterilised by UV irradiation. Human plasma fibronectin (Gibco, USA) was coated on the sterilised PMMA substrates by soaking them in 25 µg/mL fibronectin solution in Dulbecco’s phosphate buffered saline (DPBS) with calcium and magnesium (Gibco, USA) for 2 hours at 37°C. Uncoated PMMA substrates were similarly soaked in DPBS with calcium and magnesium. A small volume of the PEM suspension (5 × 10^4^ cells/substrate) was first seeded on the PMMA substrates and left to incubate for 1 hour to allow the cells to attach on the substrate. Additional fresh culture media was added to the respective cell culture wells, as required, and the PEMs were cultured for 2 days before being replated or harvested for analysis. To replate the PEMs cultured on the PMMA substrates, the cells were detached from the scaffolds with Accutase (StemCell Technologies, Canada) and seeded in respective 24-well culture plates. Replated cells were left in culture for a further 2 days before being harvested for further analysis.

#### Phalloidin- and Immuno-staining

PEMs in culture for 2 days were fixed in 10% neutral-buffered formalin solution (Sigma-Aldrich, USA) for 20 minutes and rinsed 3 times in phosphate buffered saline (PBS). Cells were permeabilised with 0.1% Triton X-100 in PBS and rinsed 3 times in PBS. The samples were blocked with 10% normal goat serum (Invitrogen, USA) for 30 minutes prior to staining.

To stain the actin filaments (F-actin), 0.16 µM Alexa Fluor 488 Phalloidin in 1% bovine serum albumin (BSA) in PBS was added; and cell nuclei were counterstained with 4’,6-Diamidino-2-Phenylindole (DAPI). Paxillin is an adaptor protein associated with focal adhesion complexes. Immunostaining with paxillin antibody was performed by incubating with 1:50 mouse anti-human paxillin (Thermo Fisher Scientific, USA) in 1% BSA in PBS, followed by Alexa Fluor 488-conjugated anti-mouse (Molecular Probes, USA) as the secondary antibody. Images were obtained with the Olympus IX71 microscope.

#### RNA isolation and cDNA synthesis

Cell lysis was carried out using TRIzolR Reagent (Life Technologies, Carlsbad, CA, USA) following manufacturer instructions. Reverse transcription of purified RNA into cDNA was performed using iScript™ cDNA synthesis kit (Bio-Rad, USA), in a reaction mix containing 1 μL iScript reverse transcriptase, 4 μL 5X iScript reaction mix and RNA template (≤ 1000 ng total RNA), made up to 20 µL with nuclease free water. The mix was incubated for 5 minutes at 25°C, 30 minutes at 42°C, 5 minutes at 85°C and holding at 4°C.

#### Real-time reverse transcriptase polymerase chain reaction (RT-PCR) analysis

RT-PCR was carried out on an iQ5 multi-colour real-time PCR detection system (Bio-Rad, USA), using the iTaq™ Universal SYBR® Green Supermix (Bio-Rad, USA). The reaction components included 5 μL iTaq™ Universal SYBR® Green Supermix, 1 μL of primer mix (5 μM forward primer, 5 μM reverse primer) and 1 to 100 ng template, made up to 10µL with nuclease free water.

The following reaction conditions were employed: 30 sec at 95°C, 45 cycles of 15 sec at 95°C and 30 sec at 60°C. Melting curve analysis was performed for fifty one 6-second cycles, with a temperature increment of 0.5°C/cycle beginning from 70°C. Target mRNA expression was calculated using the mathematical expression for fold change, 2−ΔΔC_t_, normalized to an endogenous control (GAPDH) and relative to a calibrator. A Student’s t-test was used to compare gene expression levels, where a p-value of 0.05 or less was considered statistically significant.

#### Primers design

DNA oligonucleotides were synthesized by AITbiotech (Singapore) and Integrated DNA Technologies (IDT). PCR primers for human gene expression were designed to specifically exclude pseudogenes found in the human genome. The primer sequences are listed in Table S1 and S2 (Supplementary information).

## Results

An array of methods exist for the synthesis and fabrication of topographical substrates with nanoscale features, including polymer phase separation, chemical vapor deposition, photolithography, electron beam lithography (EBL) and colloidal lithography[8]. While evaporation-induced self-assembly (EISA) has been previously used to generate crystalline substrates for other applications[9, 10], it has not been applied to create PMMA structures for cell culture, to date. In our procedure, a sintering step was carried out at a temperature slightly above the glass transition of atactic PMMA (105°C) in order to promote partial coalescence of the particles and obtain free-standing substrates of good structural integrity.

The ability of nanotopographical substrates to prevent focal adhesion assembly is governed by several guidelines, as defined by Biggs *et al*.[5]. In general, cellular adhesion is decreased on structures measuring ∼70–100 nm in height, or when the feature diameter is <70 nm. To obtain a nanoparticle substrate that satisfied the dimensional criteria, we varied final surfactant concentrations in the polymerization step to obtain different nanoparticle sizes and fabricated these into substrates by EISA. The determined particle sizes and the surface morphology of the corresponding assembled substrates are shown in Figure 1. When a final surfactant concentration 6.4 mg/mL was used, nanoparticles of average diameter 73 ± 1 nm were obtained. As these particles and the resulting assembled substrates appeared to meet the dimensional criteria for prevention of focal adhesion assembly[5], they were employed further for our cell culture experiments.

**Figure 1.**
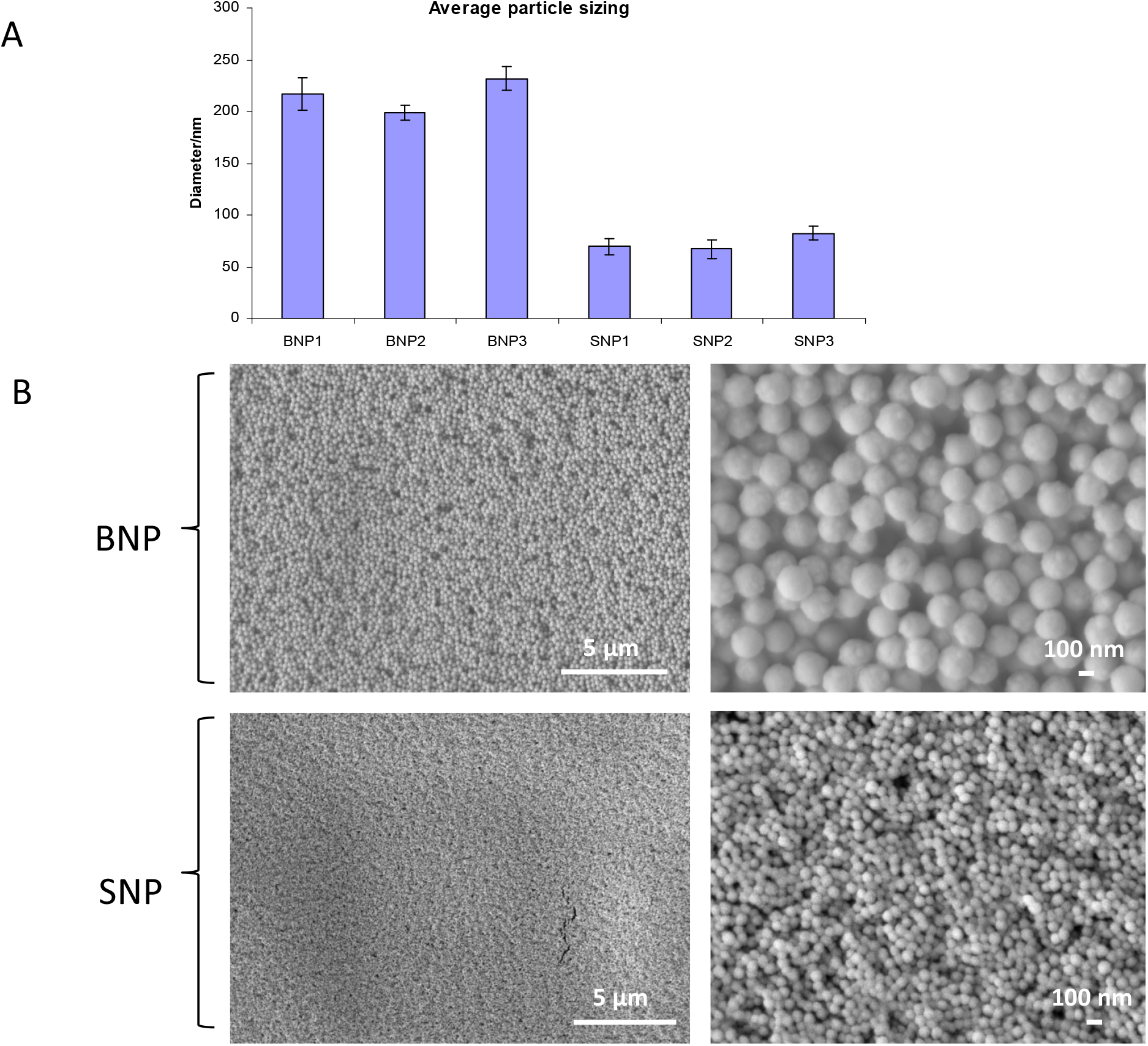
(a) Particle sizing for 3 batches of synthesized nanoparticles under conditions of lower surfactant concentration (0.60 mg/mL) and higher surfactant concentration (6.4 mg/mL), labelled BNP and SNP, respectively; (b) SEM micrographs of nanoparticle substrates assembled from the two nanoparticle types.

To verify the prevention of focal adhesion, primary melanocytes were cultured on the nanoparticle substrates for 2 days. These cells showed distinctly different morphology compared to the same cells cultured on tissue culture plate (Figure 2). While the cells cultured on the plates were of tapered morphology and relatively uniformly distributed, those on the nanostructures (both fibronectin-coated and non-coated) appeared clustered, and many of the cells were rounded. Distinct cellular foci could be observed at lower magnification, especially for the fibronectin coated substrate. The lack of cell spreading on the nanostructured substrates was indicative of restricted focal adhesion formation.

**Figure 2.**
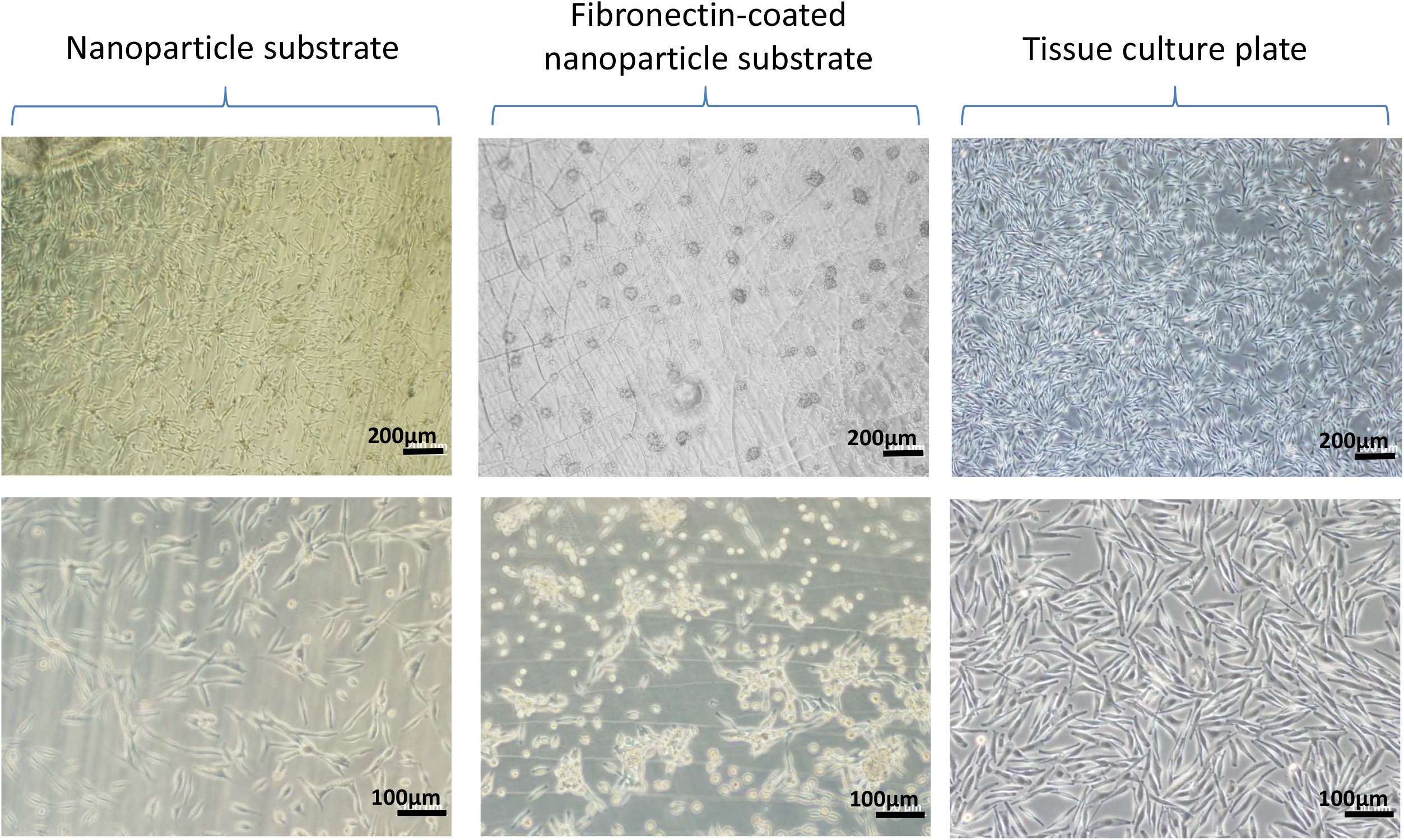
Culture of primary epidermal melanocytes on the various surfaces. Melanocytes cultured on the fibronectin-coated and non-coated nanoparticle substrates were more rounded and formed clusters.

To confirm actual prevention of focal adhesion formation, the cells were stained with paxillin, which is a focal adhesion-associated protein. Punctuate staining of paxillin was observed in the control melanocytes cultured on tissue culture plates, indicating the presence of focal adhesion complexes (Figure 3). There was a decreased expression of paxillin for the fibronectin-coated nanoparticle substrates, with almost no staining observed for the non-coated nanoparticle substrates. A similar trend was obtained for actin expression. The presence of actin stress fibers, characteristic of strong cell adhesion, was observed for melanocytes cultured on the tissue culture plates, which was reduced for the fibronectin-coated nanoparticle substrates. No actin fibers could be discerned for more rounded cells on the non-coated nanoparticles (Figure 3). An important point to note in analysing the morphology of cells on each of the substrate types was the lower number of cells on the nanoparticle substrates, both coated and non-coated. This may indicate lower viability of the cells when compared to those grown on the tissue culture plates.

**Figure 3.**
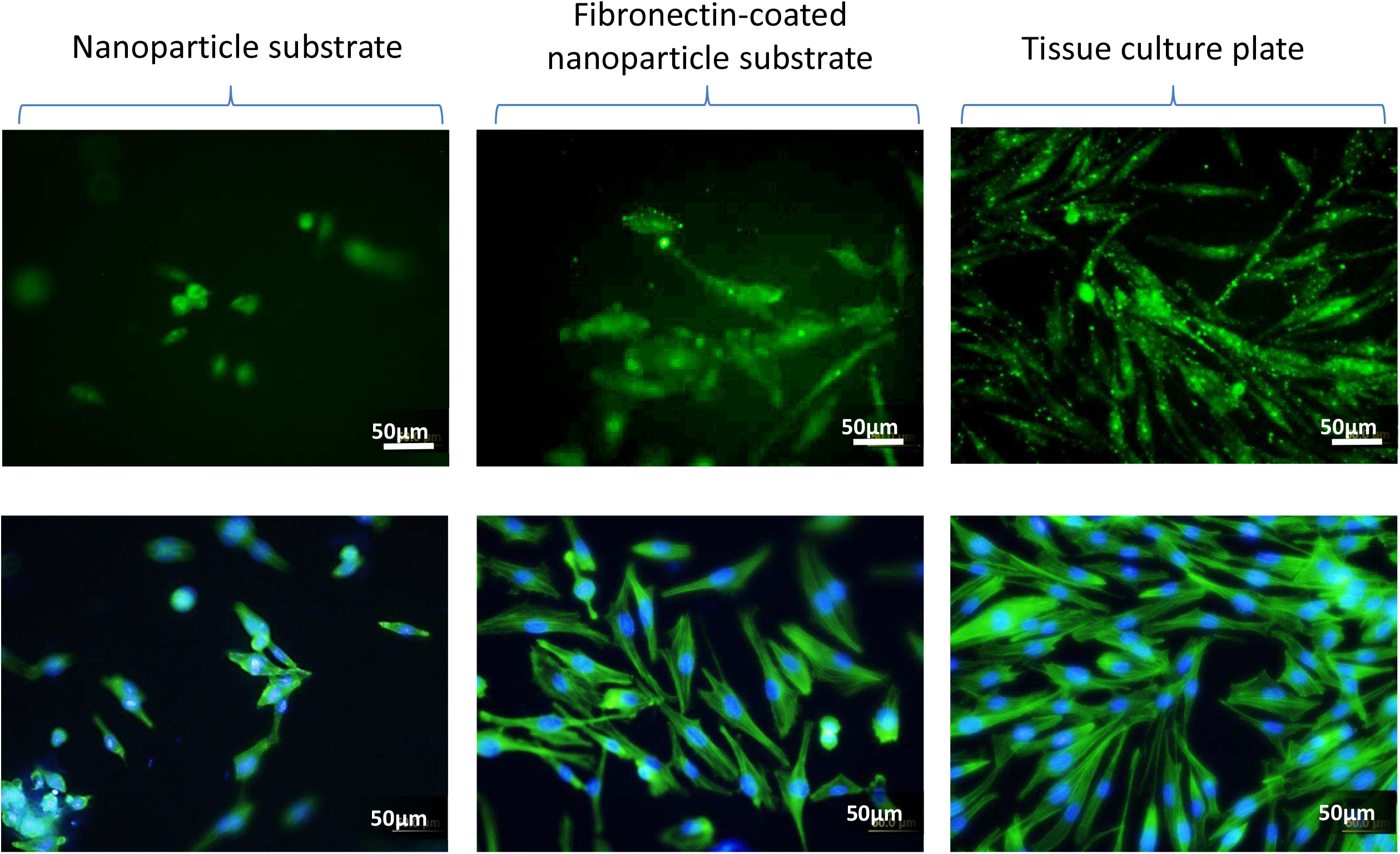
Immunostaining for paxillin (top panel) and actin (bottom panel) for PEMs cultured on the various substrates. Loss of actin stress fibers was observed for cells cultured on the non fibronectin-coated nanoparticle substrate, while cells on the fibronectin-coated substrate showed a reduction in actin staining.

Having confirmed that the cultured melanocytes had undergone a reduction in focal adhesions and were less strongly adhered to the tissue culture plates, we proceeded to quantitate the gene expression of the cells (Figure 4). Gene expression values were normalized with respect to the values obtained from culture of cells on tissue culture plastic (Relative Gene Expression of 1.0). Both fibronectin-coated and non-coated substrates exhibited increased expression (≥ 2-fold) of the epithelial markers EPCAM and KRT18, while only fibronectin-coated substrates showed increased expression of the mesenchymal markers SNAIL and TGFβ1 (Figure 4A). Interestingly, cells on both fibronectin-coated and non-coated substrates showed increased expression (≥ 2-fold) for a panel of general stemness markers comprising OCT4, NANOG, LIN28, CSPG4, NESTIN, CXCR4 and TERT, with coated substrates exhibiting higher expression of SOX2 (p < 0.05). The same trend of enhanced stemness was observed for the panel of melanocyte-specific markers (Figure 4B), with increased expression of melanocyte stemness genes and concomitant decreased expression of genes associated with the differentiated melanocyte phenotype (MITF, TYR, TRP1). The increase in stemness gene expression was more pronounced for cells cultured on the fibronectin-coated nanoparticle surfaces for the case of SOX10 gene expression (p < 0.05). Previous studies have shown that hypoxia may have a role in promoting stemness, an effect which is directly related to the expression of Hypoxia-inducible factor 1-alpha (HIF1A)[11, 12]. To exclude the possible effect of hypoxia due to the clustering of melanocytes cultured on nanoparticle substrates, the expression of the HIF1A transcript was measured and shown to be downregulated compared to cells cultured on TCP (Figure 4A).

**Figure 4.**
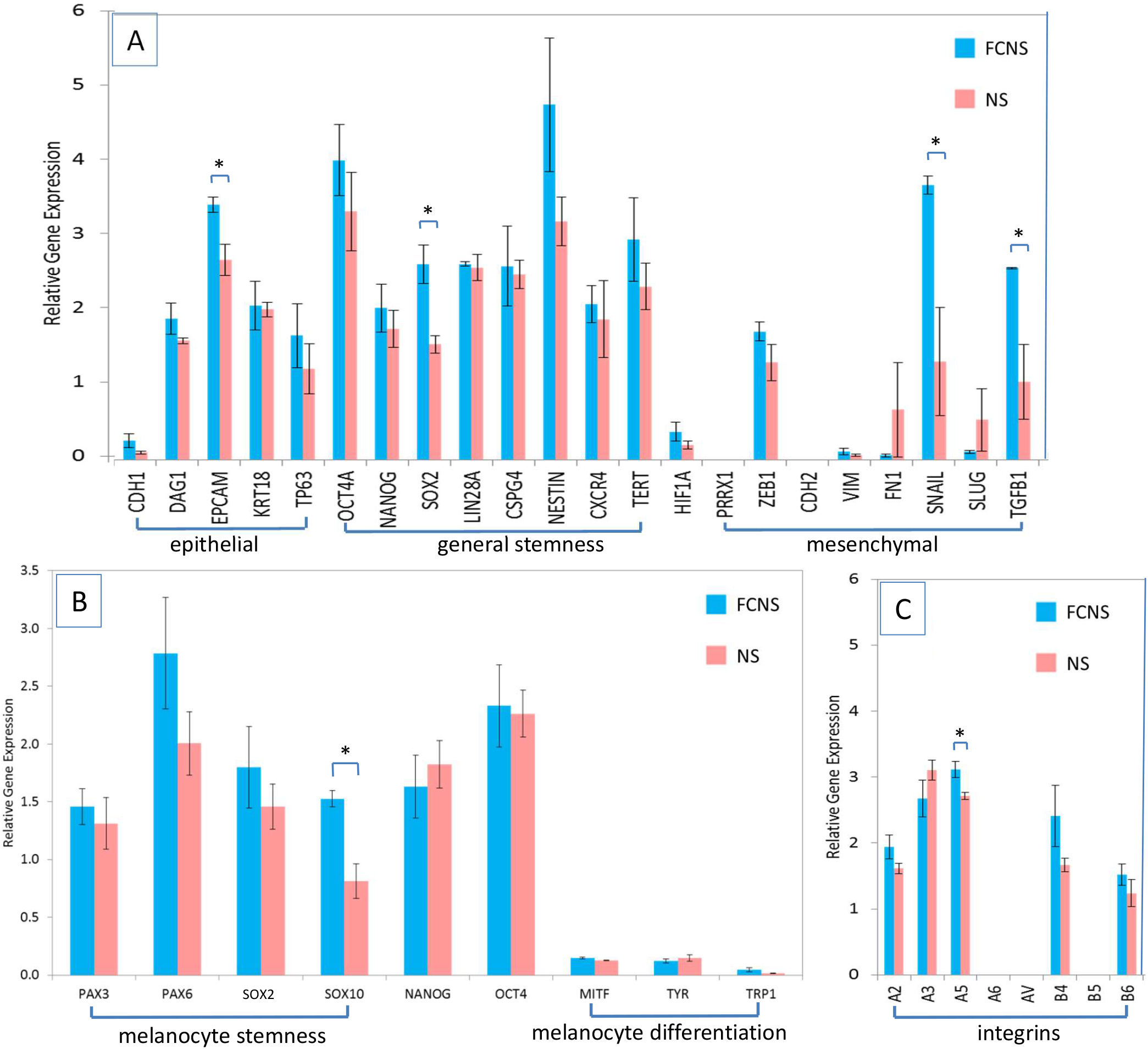
Gene expression for (A) general stemness markers, epithelial and mesenchymal markers, (B) melanocyte-specific markers, and (C) integrins, normalized with respect to expression level on tissue culture plate. Primary melanocytes were cultured on fibronectin-coated nanoparticle substrates (FCNS) and non-coated substrates (NS) (n=3). Error bars represent technical replicates. * p < 0.05; ** p <0.005.

As integrins mediate the adhesion of cells to the extracellular matrix and are involved in signalling processes that affect gene expression, we looked at the expression for a set of integrin genes. There was upregulated expression of integrins α2, α3, α5 and β4, especially for the fibronectin-coated nanoparticle substrates (Figure 4C). From our current results, integrin β6 was upregulated, while integrin β5 was not. This accords with our experimental conditions where the melanocytes were cultured on fibronectin, as it is known that integrin αVβ6 is a fibronectin receptor, whereas αVβ5 is a vitronectin receptor. As expected, integrin α6 was not upregulated as it its ligands are present mainly on laminin, but not on fibronectin.

Next, the melanocytes that had been subject to all three culture conditions were replated onto tissue culture plate). In all cases, the cells reverted to their original tapered morphologies (Figure S2). However, for the case of cells that had been cultured on the fibronectin-coated substrates prior to replating, distinct changes were observed in the gene expression of the stemness and differentiation markers after 2 days (Figure 5). Stemness markers such as PAX6, SOX10 and OCT4 were now downregulated, while the melanocyte differentiation markers were significantly upregulated (2.5-fold to 4-fold). These observations suggest that cells cultured on the fibronectin-coated nanoparticle substrates exhibited a more differentiated melanocyte phenotype when replated on tissue culture plate.

**Figure 5.**
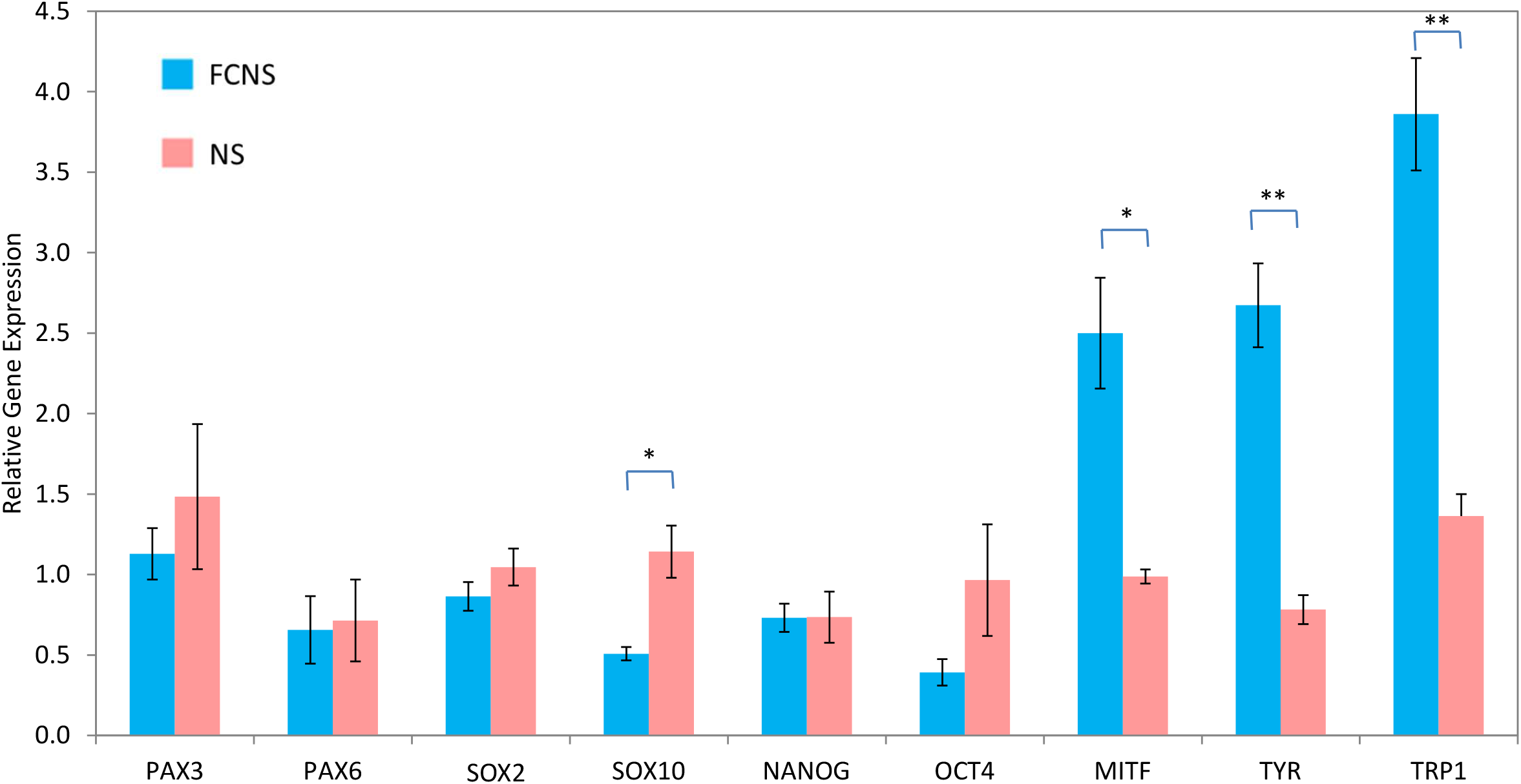
Gene expression for melanocytes that were replated onto tissue culture plate after being cultured on fibronectin-coated substrate (FCNS) and non-coated substrate (NS) for 2 days, normalized with respect to expression level on tissue culture plate (n=3). Error bars represent technical replicates. * p < 0.05; ** p <0.005.

## Discussion

### Effect of topography on focal adhesion formation

Previous work has shown that topography has a great influence on the differential gene expression and function of cells, mediated by changes in surface size features and therefore, focal adhesion arrangement[5, 13, 14]. For example, it has been demonstrated that hMSCs have an intricate sensing mechanism that allows them to differentiate between micrometer and nanosized topographies, leading to effects on cell differentiation[15]. Based on cumulated findings, general guidelines have been developed on the effect of nanofeature dimensions (portrusions or pits) on focal adhesion formation[5].

Various techniques have been used to create nanostructured surfaces for cell culture, with the objective of modulating cellular phenotype and directing cell fate[15-17]. We have employed PMMA nanoparticles synthesized by emulsion polymerisation and evaporation-induced self-assembly to obtain nanoparticle substrates that approximate the said dimensional guidelines. However, while the nanoparticles themselves are of regular dimensions, the assembled substrates of Figure 1 are not planar at the microscopic level and associated with a surface roughness consisting of random bumps and dips (Figure S1). To a lesser extent, such surface roughness is also exhibited by the tissue culture plates (TCP) used as the control surfaces in our experiment. Additionally, there may be an effect of surface chemistry (relative hydrophobicity or hydrophilicity) that contributes to the different extent of focal adhesion formation on the nanoparticle substrate, as compared to TCP. Therefore, it would be challenging to attribute the prevention of focal adhesion that is observed in our current study solely to the specific periodicity and feature size due to the assembled nanoparticles.

Nevertheless, cells cultured on the 70nm nanoparticle substrates did prevent focal adhesion formation in cultured melanocytes, as seen by the lack of paxillin and actin stress fibers. The same substrates that were coated with fibronectin also exhibited reduced focal adhesion formation, though to a lesser extent. Our subsequent experiments suggested that these reductions in focal adhesion had led to a more stem cell-like transcriptional profile compared to cells on TCP. Both fibronectin-coated and non-coated substrates showed the effect, supporting a role of topography in the differential gene expression.

Here, although the effectiveness of the fibronectin coating was not tested, a substantial coating is expected to form on PMMA under the conditions of the current coating procedure due to the high affinity of fibronectin for PMMA[18].

### Effect of focal adhesion disassembly on gene expression

Work by ourselves and others have shown that cells can revert to a phenotype associated with stem cell characteristics, when cultured on 3D scaffolds[4,19]. While cell-matrix adhesions in 3D vary depending on the physical properties and composition of the matrix, it is generally observed that in such an environment, cells show reduced formation of actin stress fibers as compared to those cultured on 2D tissue culture plates, exhibit smaller and more distributed focal adhesions, and often grow as 3D aggregates[20,21]. Under certain conditions, the phenotype of 3D-cultured cells is closely similar to that of cells that are less strongly adhered to a 2D substrate, where the formation of focal adhesions is compromised. These observations led us to hypothesize that cells cultured under conditions where focal adhesions are prevented would similarly show enhanced stemness gene expression, and this was borne out by the present experiments. Regardless of whether the nanoparticle substrates were fibronectin-coated, a significant increase was observed in the gene expression of stemness markers, accompanied by a decrease in the expression of differentiation markers.

The effect of reduced focal adhesion formation has been reported to increase the expression of stemness related genes in one other study. When mesenchymal stem cells were cultured on soft cell culture substrates, they presented with lower maturation of cell adhesions, and this was accompanied by an increase in the expression of pluripotency-related genes and a stem-like cellular phenotype[22]. These observations align well with our study, albeit their use of a different condition (soft culture substrate) to reduce focal adhesions.

For our gene expression studies, the first issue that we had to address was the development of a panel of markers to measure changes in the stemness of the melanocytes. PAX3 and SOX10 have been identified as two of the genes expressed by melanocyte stem cells and were used as a measure of stemness.[23-27]. Besides these, we measured the expression of PAX6 and SOX2, which are neuroectodermal markers and thus expressed by melanocyte precursors[]. In addition, we also measured the levels of general stemness markers expressed by pluripotent/somatic stem cells (such as OCT4, NANOG, LIN28A, CSPG4, TERT, CXCR4) and those that are expressed by differentiated melanocytes and associated with their function, namely MITF, TYR and TRP1[9]. Examining the pattern of gene expression changes allowed us to ascertain whether the melanocytes were proceeding towards, or away from a stem cell like-phenotype.

### ECM-coating of nanoparticle substrate

Fibronectin was chosen as the extracellular matrix (ECM) molecule to coat the nanoparticle substrates as it has been shown to play a role in melanocyte adhesion and viability [31]. Fibronectin harbours the ligands for integrin α5β1, which is a primary receptor for attachment of melanocytes [30, 31]. Changes in cell shape and dimensionality has been reported to influence the use of specific integrins in cell adhesions[32] as well as the relative gene and protein expression levels of the various integrins[4]. The upregulated expression of integrin α5β1, as observed in the present study, may lead to further ECM-integrin binding and signalling for the fibronectin-coated nanoparticle substrate, and this is postulated to result in the higher upregulation of stemness markers for the latter. From our experiments, this led to a more differentiated melanocyte phenotype when the cells were replated on tissue culture plate. From the viewpoint of clinical practice, our experiments suggest the potential of modulating cell culture conditions to obtain more functionally differentiated melanocytes for transplantation.

## Conclusion

Nanostructured substrates obtained by evaporation-induced self-assembly of emulsion polymerized PMMA nanoparticles offer excellent surfaces for the culture of cells to prevent, or reduce, focal adhesion formation. For the case of melanocytes, results from our present study suggest that (1) prevention of focal adhesion formation is associated with a more stem cell-like transcriptional profile, and (2) fibronectin-coating promotes the formation of an intermediate adhesive state with higher differentiation potential.

## Acknowledgment

Funding was provided by the Institute of Bioengineering and Nanotechnology (Biomedical Research Council, Agency for Science, Technology and Research, Singapore).

## REFERENCES

[1] Burridge K, Guilluy C. Focal adhesions, stress fibers and mechanical tension. Experimental cell research. 2016;343:14–20.

[2] Murphy-Ullrich JE, Sage EH. Revisiting the matricellular concept. Matrix biology : journal of the International Society for Matrix Biology. 2014;37:1–14.

[3] Murphy-Ullrich JE. The de-adhesive activity of matricellular proteins: is intermediate cell adhesion an adaptive state? The Journal of clinical investigation. 2001;107:785–90.

[4] Ma NK, Lim JK, Leong MF, Sandanaraj E, Ang BT, Tang C, et al. Collaboration of 3D context and extracellular matrix in the development of glioma stemness in a 3D model. Biomaterials. 2016;78:62–73.

[5] Biggs MJ, Richards RG, Dalby MJ. Nanotopographical modification: a regulator of cellular function through focal adhesions. Nanomedicine : nanotechnology, biology, and medicine. 2010;6:619–33.

[6] Yuan L, Wang Y, Pan M, Rempel GL, Pan Q. Synthesis of poly(methyl methacrylate) nanoparticles via differential microemulsion polymerization. European Polymer Journal. 2013;49:41–8.

[7] Sailaja AK, Amareshwar P, Dakshayani M. Preparation of polymethylmethacrylate nanoparticles by emulsion polymerization technique. Asian Journal of Biochemical and Pharmaceutical Research. 2011;1:298–302.

[8] Norman JJ, Desai TA. Methods for fabrication of nanoscale topography for tissue engineering scaffolds. Annals of biomedical engineering. 2006;34:89–101.

[9] Brezesinsky T, Groenewolt M. A. G, Pinna N, Antonietti M, Smarsly BM. Evaporation-Induced Self-Assembly (EISA) at Its Limit: Ultrathin, Crystalline Patterns by Templating of Micellar Monolayers. 2006;18:2260–3.

[10] Nam H, Jung D, Yi G, Choi H. Close-Packed Hemispherical Microlens Array from Two-Dimensional Ordered Polymeric Microspheres. Langmuir. 2006;22:7358–63.

[11] Yun Z, Lin Q. Hypoxia and regulation of cancer cell stemness. Advances in experimental medicine and biology. 2014;772:41–53.

[12] Liu Y, Mukhopadhyay P, Pisano MM, Lu X, Huang L, Lu Q, et al. Repression of Zeb1 and hypoxia cause sequential mesenchymal-to-epithelial transition and induction of aid, Oct4, and Dnmt1, leading to immortalization and multipotential reprogramming of fibroblasts in spheres. Stem cells. 2013;31:1350–62.

[13] M.J. Dalby, N. Gadegaard, R. Tare, A. Andar, M.O. Riehle, P. Herzyk, C.D. Wilkinson, R.O. Oreffo, The control of human mesenchymal cell differentiation using nanoscale symmetry and disorder, Nat Mater 6(12) (2007) 997–1003.

[14] B.K. Teo, S.T. Wong, C.K. Lim, T.Y. Kung, C.H. Yap, Y. Ramagopal, L.H. Romer, E.K. Yim, Nanotopography modulates mechanotransduction of stem cells and induces differentiation through focal adhesion kinase, ACS Nano 7(6) (2013) 4785–98.

[15] B.K. Teo, S. Ankam, L.Y. Chan, E.K. Yim, Nanotopography/mechanical induction of stem-cell differentiation, Methods Cell Biol 98 (2010) 241–94.

[16] Dalby MJ, Gadegaard N, Oreffo RO. Harnessing nanotopography and integrin-matrix interactions to influence stem cell fate. Nature materials. 2014;13:558–69.

[17] H. Yu, Y.S. Lui, S. Xiong, W.S. Leong, F. Wen, H. Nurkahfianto, S. Rana, D.T. Leong, K.W. Ng, L.P. Tan, Insights into the role of focal adhesion modulation in myogenic differentiation of human mesenchymal stem cells, Stem Cells Dev 22(1) (2013) 136–47.

[18] P.E. Vaudaux, F.A. Waldvogel, J.J. Morgenthaler, U.E. Nydegger, Adsorption of fibronectin onto polymethylmethacrylate and promotion of Staphylococcus aureus adherence, Infect Immun 45(3) (1984) 768–74.

[19] Wang K, Kievit FM, Erickson AE, Silber JR, Ellenbogen RG, Zhang M. Culture on 3D Chitosan-Hyaluronic Acid Scaffolds Enhances Stem Cell Marker Expression and Drug Resistance in Human Glioblastoma Cancer Stem Cells. Advanced healthcare materials. 2016;5:3173–81.

[20] Edmondson R, Broglie JJ, Adcock AF, Yang L. Three-dimensional cell culture systems and their applications in drug discovery and cell-based biosensors. Assay and drug development technologies. 2014;12:207–18.

[21] Harunaga JS, Yamada KM. Cell-matrix adhesions in 3D. Matrix biology : journal of the International Society for Matrix Biology. 2011;30:363–8.

[22] H. Gerardo, A. Lima, J. Carvalho, J.R.D. Ramos, S. Couceiro, R.D.M. Travasso, R. Pires das Neves, M. Grãos, Soft culture substrates favor stem-like cellular phenotype and facilitate reprogramming of human mesenchymal stem/stromal cells (hMSCs) through mechanotransduction, Sci Rep 9(1) (2019) 9086.

[23] Kubic JD, Young KP, Plummer RS, Ludvik AE, Lang D. Pigmentation PAX-ways: the role of Pax3 in melanogenesis, melanocyte stem cell maintenance, and disease. Pigment cell & melanoma research. 2008;21:627–45.

[24] Lang D, Lu MM, Huang L, Engleka KA, Zhang M, Chu EY, et al. Pax3 functions at a nodal point in melanocyte stem cell differentiation. Nature. 2005;433:884–7.

[25] Harris ML, Buac K, Shakhova O, Hakami RM, Wegner M, Sommer L, et al. A dual role for SOX10 in the maintenance of the postnatal melanocyte lineage and the differentiation of melanocyte stem cell progenitors. PLoS genetics. 2013;9:e1003644.

[26] Osawa M, Egawa G, Mak SS, Moriyama M, Freter R, Yonetani S, et al. Molecular characterization of melanocyte stem cells in their niche. Development. 2005;132:5589–99.

[27] Li A. The biology of melanocyte and melanocyte stem cell. Acta biochimica et biophysica Sinica. 2014;46:255–60.

[28] Zhang X, Huang CT, Chen J, Pankratz MT, Xi J, Li J, et al. Pax6 is a human neuroectoderm cell fate determinant. Cell stem cell. 2010;7:90–100.

[29] Hosaka C, Kunisada M, Koyanagi-Aoi M, Masaki T, Takemori C, Taniguchi-Ikeda M, et al. Induced pluripotent stem cell-derived melanocyte precursor cells undergoing differentiation into melanocytes. Pigment cell & melanoma research. 2019.

[30] Beliveau A, Berube M, Rousseau A, Pelletier G, Guerin SL. Expression of integrin alpha5beta1 and MMPs associated with epithelioid morphology and malignancy of uveal melanoma. Investigative ophthalmology & visual science. 2000;41:2363–72.

[31] Scott G, Cassidy L, Busacco A. Fibronectin suppresses apoptosis in normal human melanocytes through an integrin-dependent mechanism. J Invest Dermatol. 1997 Feb;108(2):147–53. doi: 10.1111/1523-1747.ep12332650. PMID: 9008226.

[32] Cukierman E, Pankov R, Stevens DR, Yamada KM. Taking cell-matrix adhesions to the third dimension. Science. 2001;294:1708–12.

