## Supplementary material for "Focal adhesion prevention on nanoparticle substrates upregulates stemness gene expression in primary melanocytes": SI

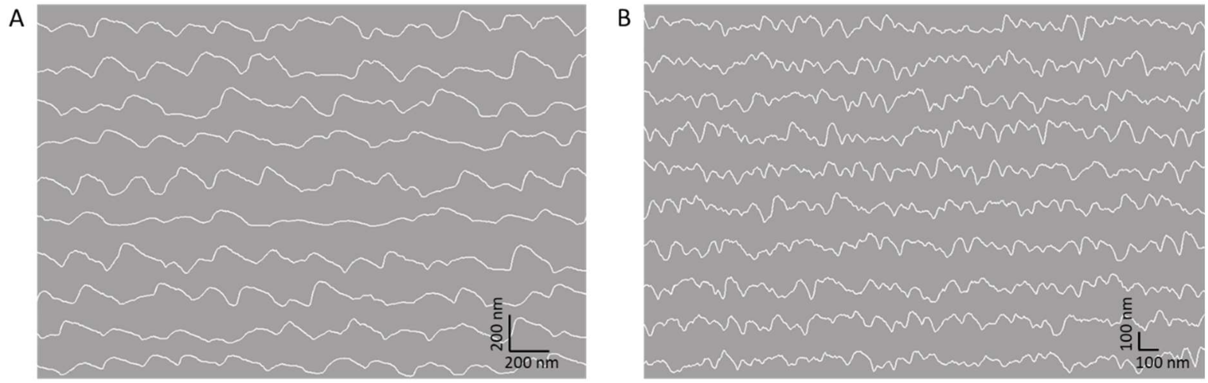


Figure S1. Surface profile of (A) BNP and (B) SNP, by image analysis of substrates assembled using the two particle sizes in Figure 1, where the height of each line profile corresponds to the grayscale intensity of the image. The periodicity of the surface, corresponding to each particle size, is observed.

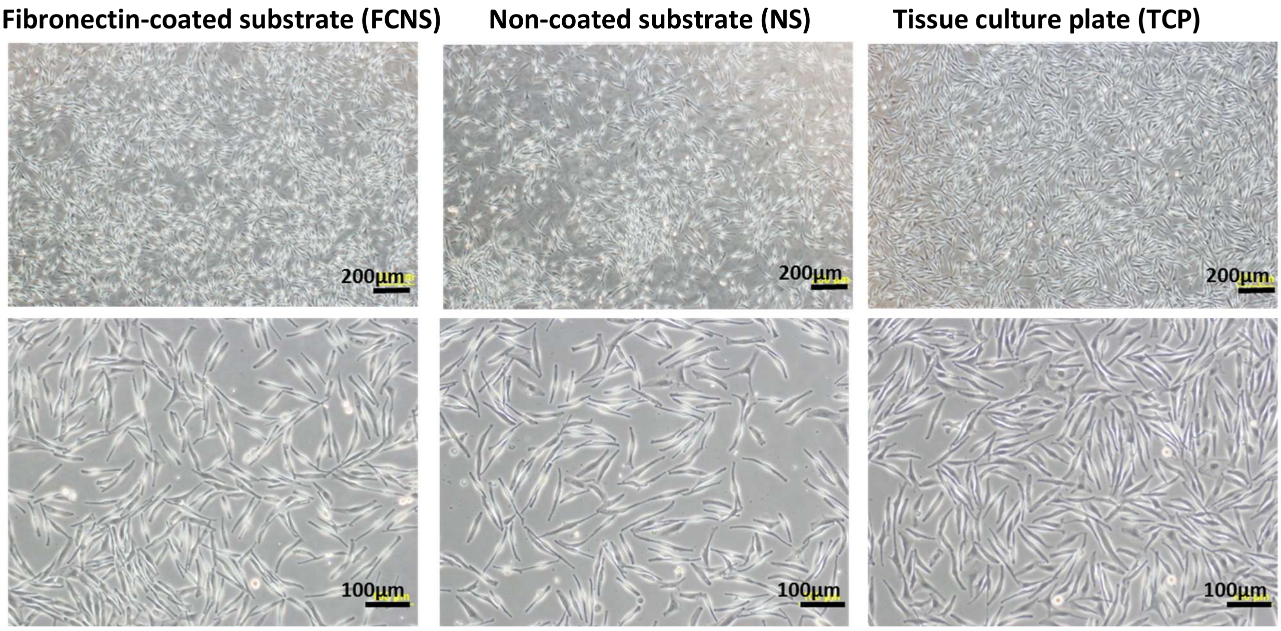


Figure S2. After replating on tissue culture plate, melanocytes cultured under all three conditions revert to tapered morphology.

**Supplementary Tables**

Table S1. Real-time PCR primer sequences for general stemness markers and integrins (Fig 4A,C)

| Target | Forward primer (5’ - 3') | Reverse primer (5' - 3') |
|---|---|---|
| GAPDH | TTGACGCTGGGGCTGGCATT | GTGCTCTTGCTGGGGCTGGT |
| ITG A2 | CCTACAATGTTGGTCTCCCAGA | AGTAACCAGTTGCCTTTTGGATT |
| ITG A3 | TCAACCTGGATACCCGATTCC | GCTCTGTCTGCCGATGGAG |
| ITG A5 | TCGTGTCCGCTAGTGCCTCC | GATGCAGGCCACAGGGTTCC |
| ITG A6 | ATGCACGCGGATCGAGTTT | TTCCTGCTTCGTATTAACATGCT |
| ITG AV | ATCTGTGAGGTCGAAACAGGA | TGGAGCATACTCAACAGTCTTTG |
| ITG B4 | AGCAGACCAAGTTCCGGCAGCA | GCGCCATCAGCACTGTGTCCAC |
| OCT4A | TGGAGAAGGAGAAGCTGGAGCAAAA | GGCAGATGGTCGTTTGGCTGAATA |
| NANOG | AATACCTCAGCCTCCAGCAGATG | TGCGTCACACCATTGCTATTCTTC |
| SOX2 | GCGGGGGAATGGACCTTGTA | TTCCTGCAAAGCTCCTACCGT |
| LIN28A | GAGCATGCAGAAGCGCAGATCAAA | TATGGCTGATGCTCTGGCAGAAGT |
| LIN28B | ATATCGGTGTGCTGTGATGC | TGAGCAACGCTTATCATGTTTT |
| CSPG4 | CTGTGGTGCTGACTGTCGTAGA | GGTAGGGCAGGCCAAGGGTC |
| NESTIN | CTTCCCTCCGCATCCCGTCA | AAAGCCAGCATGTCACCCTCC |
| CXCR4 | ACGGACAAGTACAGGCTGCAC | CCAGAAGGGAAGCGTGATGACA |
| CDH1 | ACGCTGTGTCATCCAACGGG | CCTCCTGGGTGAATTCGGGCTT |
| DAG1 | CTCTCTGTGGTTATGGCTCAGT | CTGTTGGAATGGTCACTCGAAAT |
| EPCAM | GGACCTGACAGTAAATGGGGAACA | ACAACTGCTATCACCACAACCACA |
| KRT18 | GGCATCCAGAACGAGAAGGAG | ATTGTCCACAGTATTTGCGAAGA |
| PRRX1 | CTGATGCTTTTGTGCGAGAA | ACTTGGCTCTTCGGTTCTGA |
| ZEB1 | GGGCGACCAAGAACAGGACT | GTGTGGGACTGCCTGGTGAT |
| CDH2 | GGTGGAGGAGAAGAAGACCAGG | GGCATCAGGCTCCACAGTGT |
| VIM | CCTTGAACGCAAAGTGGAATC | GACATGCTGTTCCTGAATCTGAG |
| FN1 | TACTGGCCTGGAACCGGGAA | ACCAGTTGGGGAAGCTCGTC |
| Snail | ACGGCCTAGCGAGTGGTTCT | GATTGGGGTCGGAGGGCTTC |
| Slug | AGCTTTCAGACCCCCATGCC | TGGCCAGCCCAGAAAAAGTTGA |
| TGF-β1 | GGGCAGATCCTGTCCAAGC | GTGGGTTTCCACCATTAGCAC |
| HIF1A | GAAAGCGCAAGTCCTCAAAG | TGGGTAGGAGATGGAGATGC |
| TERT | AACCTTCCTCAGCTATGCCCG | CAGCCGCAAGACCCCAAAGA |
| TP63 | CCAAAGCGAGGCACCCTTA | GGAGAGTAGGCTGCCATGAGG |

Table S2. Real-time PCR primer sequences for melanocyte stemness and differentiation markers (Figs 4B, 5)

| Target | Forward primer (5’ - 3') | Reverse primer (5' - 3') |
|---|---|---|
| NANOG | CAAAGGCAAACAACCCACTT | TCTGCTGGAGGCTGAGGTAT |
| OCT4 | CTTGCTGCAGAAGTGGGTGGAGGAA | CTGCAGTGTGGGTTTCGGGCA |
| SOX2 | TCGGCATCGCGGTTTTT | ACAGCAAATGACAGCTGCAAA |
| SOX10 | AGCCCAGGTGAAGACAGAGA | ATAGGGTCCTGAGGGCTG AT |
| PAX3 | CTGGAACATTTGCCCAGACT | GCTGTCGGTTCCTAGTCCAG |
| PAX6 | GCCAGCAACACACCTAGTCA | TGTGAGGGCTGTGTCTGTTC |
| MITF | CCGGGTGCAGAATTGTAACT | GGACAATTTTGGCATTTTGG |
| TRP1 | AGCAGTAGTTGGCGCTTTGT | TCAGTGAGGAGAGGCTGGTT |
| TYR | TTGTACTGCCTGCTGTGGAG | CAGGAACCTCTGCCTGAAAG |

GAPDH: Glyceraldehyde 3-phosphate dehydrogenase, ITG: Integrin, OCT4A: Octamer-binding transcription factor 4A, NANOG: homeobox transcription factor Nanog, SOX2: SRY (sex determining region Y)- box 2, LIN28A: lin-28 homolog A, CSPG4: Chondroitin sulfate proteoglycan 4, CXCR4: Chemokine (C-X-C motif) receptor 4, KRT18: Keratin, type I cytoskeletal 18, PRRX1: Paired related homeobox 1, ZEB1: Zinc finger E-box-binding homeobox 1, CDH1: Cadherin-1, CDH2: Cadherin-2, EPCAM: Epithelial cell adhesion molecule, DAG1: dystroglycan 1, VIM: vimentin, FN1: fibronectin 1, Snail: Zinc finger protein SNAI1, Slug: Zinc finger protein SNAI2, TGF- β1: Transforming growth factor beta 1, HIF1A: Hypoxia-inducible factor 1-alpha, TERT: Telomerase reverse transcriptase, TP63: tumor protein p63, SOX10: SRY (sex determining region Y)- box 10, PAX3: paired box gene 3, PAX6: paired box gene 6, MITF: melanocyte inducing transcription factor, TRP1: Tyrosinase related protein 1, TYR: tyrosinase.
